# Active microrheology using pulsed optical tweezers to probe viscoelasticity of Lamin A towards diagnosis of laminopathies

**DOI:** 10.1101/2021.02.05.429901

**Authors:** C. Mukherjee, A. Kundu, R. Dey, A. Banerjee, K. Sengupta

**Author notes:** Authors contributed equally (C. Mukherjee, A. Kundu).

## Abstract

Lamins are nucleoskeletal proteins of mammalian cells that stabilize the structure and maintain the rigidity of the nucleus. These type V intermediate filament proteins which are predominantly of A and B types provide necessary tensile strength to the nucleus. Single amino acid missense mutations occurring all over the lamin A protein form a cluster of human diseases termed as laminopathies, a few of which principally affect the muscle and cardiac tissues responsible for load bearing functionalities of the body. One such mutation is lamin A350P which causes dilated cardiomyopathy in patients. It is likely that a change from alanine to proline in the α-helical 2B rod domain of the protein might severely disrupt the propensity of the filaments to polymerise into functional higher order structures required to form a fully functional lamina with its characteristic elasticity. In this study, we validate for the very first time, the application of active microrheology employing oscillating optical tweezers to investigate any alterations in the visco-elastic parameters of the mutant protein meshwork *in vitro*, which might translate into possible changes in nuclear plasticity. We confirm our findings from this robust yet fast method by imaging both the wild type and mutant lamins using a super resolution microscope, and observe changes in the mesh size which explain our measured changes in the viscoelastic parameters of the lamins. This method could naturally be extended to conduct microrheological measurements on any intermediate filament protein or any protein endowed with elastic behavior, with minor schematic modifications, thus bearing significant implications in laminopathies and other diseases which are associated with changes in structural rigidity of any cellular organelle.

**Significance:** Lamin A mutations produce an array of diseases termed as laminopathies which are primarily characterized by alteration of elastic behavior of the nucleus which in turn leads to defects in mechanotransduction. This is the first report in the lamin arena which shows a fast, accurate and direct quantification of elastic moduli of lamin A using optical tweezers-based microrheology. This has very significant implications and can be registered to be a robust and universal method that could also be suitably used for probing changes in elastic properties of any proteins or surfactants in a disease scenario such as SARS-Cov2 (Covid-19), which is pandemic at this time.

## Introduction

Nuclear lamin proteins form a viscoelastic meshwork which act as a scaffold for the foundation of the nuclear membrane and thereby build a framework for the nuclear entity in all metazoan cells [1]. Apart from rendering stiffness to the nucleus, lamins are also involved in the organization of chromatins, replication, DNA damage repair, transcription, and positioning of nuclear pore complex (NPC) [1, 2]. The 14 nm thick filamentous layer of mostly insoluble type V intermediate filament (IF) proteins is present in close apposition to the inner nuclear membrane (INM) called the nuclear lamina, while a fraction of unpolymerized soluble protein is ubiquitously distributed throughout the nucleoplasm [3, 4]. While the laminar fraction imparts tensile strength, the nucleoplasmic fraction highlights viscoelastic character of the nucleus [5]. The location and solubility of lamins depend upon phosphorylation pattern and also affect its function at different stages of the cell cycle [1, 6]. The nuclear lamina is occupied by two types of relatively identical protein fibres namely lamin A (A and C) and B (B1 and B2) varying in molecular weights and post translational modifications. Lamin A/C arises from alternative splicing of *LMNA* gene whereas B1 and B2 proteins are product of *LMNB1* and *LMNB2* genes respectively [1]. While lamin B proteins are expressed in all cell types from embryonic stage throughout development, lamin A is developmentally regulated and expressed in differentiated mammalian somatic cells, including cardiomyocytes [2, 7]. From the homotypic as well as heterotypic interaction observed between purified lamin A and B *in-vitro* and *ex-vivo*, it is believed that they form both interconnected as well as markedly distinct networks inside the nucleus [8-10]. When lamin A/C was knocked down in human epithelial cells, the high deformability of the nucleus was comparable to the fluid like adult haematopoetic stem cells having only lamin B, thus indicating the importance of lamin A:B stoichiometry and also establishing the role of lamin A alone in the regulation of nuclear viscoelasticity [5, 11, 12]. Like all IF proteins, lamins also share a structurally conserved helical rod domain flanked by a variable unstructured N terminal head and a globular C terminal tail region [13-17]. The primarily α-helical rod domain of the 664 amino acid long pre-lamin A protein is composed of 350 amino acid residues which can be further subdivided into 1A, 1B, 2A and 2B domains with heptad repeat periodicity (Figure 1A) in each of the domains. Apart from forming the filament backbone, the rod domains mainly impart tensile properties to the protein by participating in self-assembly and formation of a coiled coil dimer to form higher order structures [14, 18]. Recently our group established the roles of 1B and 2B in modulating the elasticity of lamin A [19]. The dimers are composed of parallel and unstaggered monomers, which can further arrange themselves *in-vitro* in a polar head to tail fashion involving the N and C terminal end segments. By using the SILAC cross-linking mass spectroscopy, it was recently demonstrated that the coiled coil dimers slide past each other in a spring like contraction contributing to the flexibility properties of the lamin polymer which can be disrupted by disease causing mutations in the protein [18]. While mutations associated with *LMNB1 and LMNB2* genes are embryonically lethal saving for a few exceptions, lamin A has been found to be associated with around 500 single amino acid substitutions throughout the length of the protein leading to 16 distinct human diseases termed as Laminopathies [18, 20] (http://www.umd.be/LMNA/). Dilated cardiomyopathy (DCM) is amongst one of the most commonly occurring autosomal dominant form (associated with more than 145 lamin A mutations) of laminopathies causing myocardial disorders in patients. It is characterized by early atrial fibrillation, left ventricular non-compaction, right ventricular arrhythmia, conduction defect, even causing death in severe conditions [20, 21]. Evidence shows that *LMNA* gene mutations give rise to mechanically weaker and misshapen nuclei with frequent breakage of the nuclear envelope, thereby exposing the chromatin, and eventually culminating into cell death [22, 23]. It can be concluded from phenotypic observations that most LMNA mutations specifically affect the tissue cells which are mechanically resilient like myocytes, adipocytes and osteocytes. Hence it is not surprising that even though lamin A is present in all tissue types, it’s expression from brain to bones scales with the tissue stiffness and so does the disease progression [11, 18].

**Figure 1.**
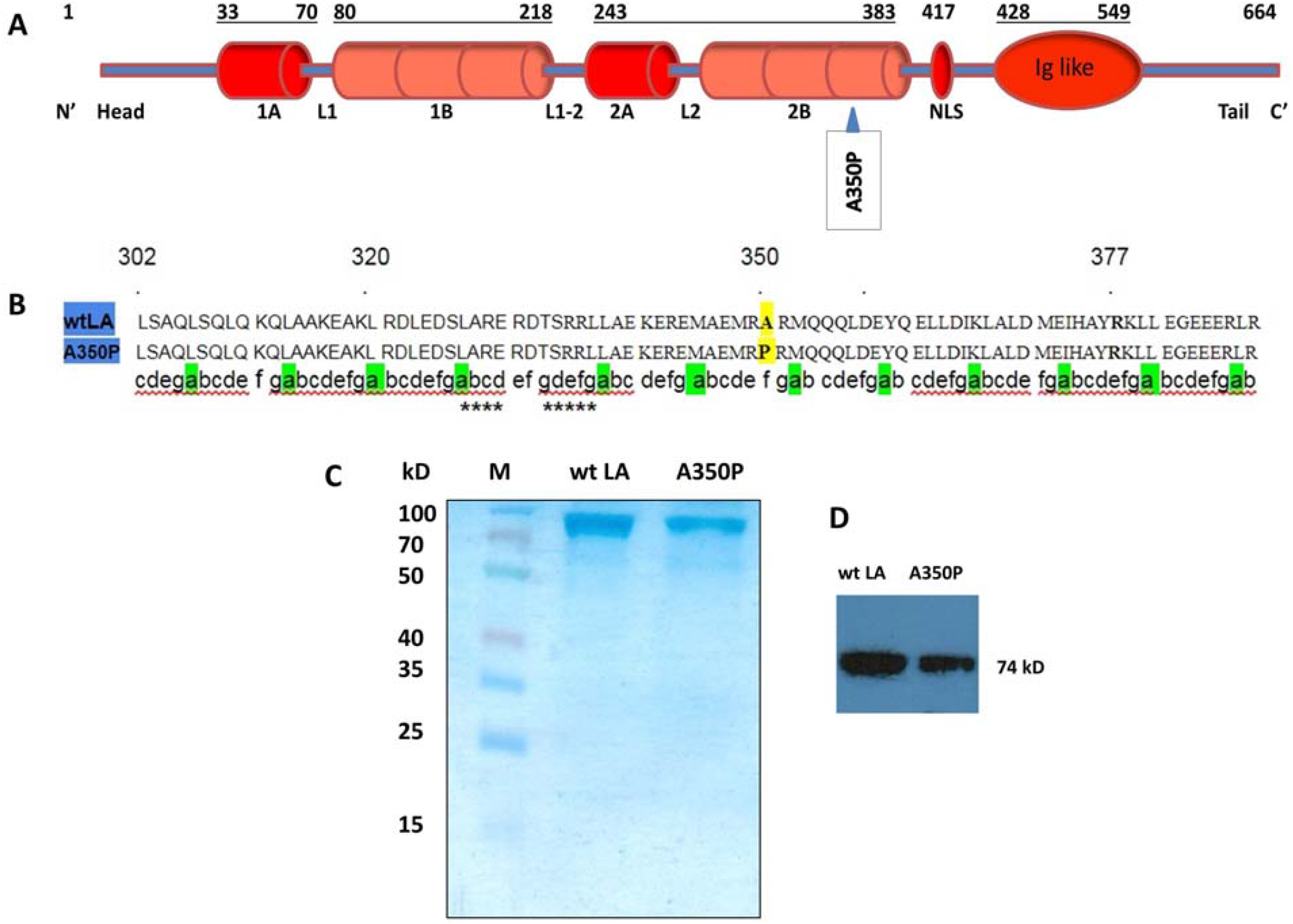
Lamin A proteins and their purification. (A) Schematic of the full length lamin A protein showing the region of mutation. (B) Amino acids of the 2B domain of the lamin A protein form α-helical coiled coil and shows their signature heptad repeat. In comparison to wt LA, the A350P mutant has a single amino acid mutation turning alanine at the 350^th^ position to proline. (C) 10% SDS PAGE analysis of pure fractions of wt LA, and A350P from Resource Q column. Numbers corresponding to the bands of the marker in lane M are in kilo Daltons. (D) Immunoblot of the same fractions using mouse monoclonal anti lamin A antibody.

Cell stretching assays and microfluidic platforms have confirmed that cells from tissues expressing LMNA mutants in drosophila, mouse and human samples have readily deformable nuclei compared to the healthy controls with concomitant defects in force transmission between nucleus and cytoskeleton [24-26]. Earlier data from our laboratory elucidated significant alterations in the secondary and tertiary structures and enthalpy changes in the deoligomerisation process of human lamin A mutants as well as aberrant laminar organization in mammalian cells expressing the mutants associated with DCM [21, 27]. The pathophysiology of the disease can be extended to the change in local network anisotropies of the mutant proteins, which induces a change in the viscoelastic property of the ensemble network [28]. In retrospect, previous data on the quantitative rheological measurements of the full length lamin A *in vitro* delineated a stiffening property of the differential elastic modulus on superposition of rotation and oscillatory shear stress for DCM specific mutants [28]. Also, single molecule force spectroscopic studies revealed altered mechanoresistant properties of lamin A tail domains specific for EDMD mutants [29].

Optical tweezers, due to their ability to confine and manipulate single mesoscopic particles in a fluidic medium, have facilitated both active and passive microrheology in a large range of viscous and viscoelastic fluids including soft biological materials [30]. Now, we have recently developed a new technique of fast active microrheology, where we use pulse-scanned optical tweezers to measure the response of a probe particle in a viscoelastic medium, and thereby extract the frequency dependent complex shear modulus *G*(ω) [30] over a large frequency range. In the present study, we have used this methodology to compare the viscoelastic properties of homologous polymers of wild type lamin A and its mutant A350P protein (henceforth referred to as wt LA and A350P respectively) *in vitro* at varying concentrations, and observe how the parameters evolve with frequency. It may be noted that the mutant gene variant was initially reported in 4 Russian patients identified with DCM. 2 of the 4 probands were diagnosed with the myopathy while they were as young as 14 and 19. Three of them were diagnosed with premature ventricular contractions (PVC) and atrial fibrillation while two had left bundle branch block (LBBB) and atrioventricular blockage. The only female amongst the probands along with another male patient had pacemakers installed as a treatment to their arrhythmias [31]. The mutation is positioned at exon 6 of LMNA gene which codes for most of the 2B rod domain of the protein (http://www.umd.be/LMNA/). Since proline has been well documented to be an α-helix destabilizer because of the irregular geometry of its structure which forces a bend or a kink in the helix [32], the importance of the study of this particular mutant variant is understandable. In our experiments we obtained a large characteristic difference in terms of compliance between the mutant and the wild type. A350P polymers showed more elastic hardening than the wt LA. Our conclusions were confirmed by super resolution microscope imaging of both lamins – which revealed the mesh size of the polymer network of the mutant to be considerably smaller and compact compared to the wild type – thus accounting for the greater elastic hardening of the former.

## Materials and Methods

### Site directed mutagenesis (SDM) and cloning

The mutant A350P mutant was generated using pET-LA as template by site directed mutagenesis (QuickChange II XL Site Directed Mutagenesis Kit, Agilent-USA). The primers used for the same were a) A350P forward 5’-gctgcatccggccgcatctcggc-3’ and b) A350P reverse 5’-gccgagatgcggccaaggatgcagc-3’ (Sigma Aldrich, India). Full length A350P was thereafter cloned into an empty pEGFP (Clontech, USA) vector using the following amplification primers: a) forward: 5’-ccggaattccggatggagaccccgtcccag-3’ and b) reverse: 5’-cggggtaccccgttacatgatgctgcagttctgggggc-3’. The restriction enzymes used for the same were EcoR1-HF and Kpn1-HF respectively (NEB, USA). SDM and cloning were performed as described earlier [21, 27, 28] Sanger sequencing was employed in each case to confirm the mutation and correct insertion (Eurofins Genomics, India).

### Protein expression and purification

Wild type and A350P full length lamin A proteins were expressed from pET-28a(+) by transformation into BL21(DE3)pLysS competent cells. Positive colonies were induced with 1mM IPTG (Thermo Scientific, USA), at OD_600_ of 0.5 in 2xYT medium (Tryptone and NaCl from Himedia, India) for 5 hours in the presence of Kanamycin and Chloramphenicol (USB corporation, USA) at 37°C for protein expression as described earlier [21, 27, 28]. Cell lysate was prepared as described by Moir *et*.*al*. 1991 [33]. Proteins were purified to near homogeneity over consecutive running on MonoQ ion exchange column and Superdex 200 increase 10/300 GL pre-packed Tricorn column (GE Healthcare, Sweden) in 6 M urea, 25mM Tris-HCl (pH 8.5), 250 mM NaCl and 1 mM DTT (Sigma Aldrich,USA) as described earlier [21, 27, 28]. Proteins were renatured by dialyzing out urea in a step wise manner from 6 M in steps of two at room temperature using Slide–A-Lyzer MINI Dialysis units (Thermo Scientific, USA) with a 10,000 Dalton MWCO. All the experiments have been conducted in the working buffer (25 mM Tris-HCl pH 8.5, 250 mM NaCl and 1 mM DTT) for both the wt LA and A350P proteins. Proteins were resolved on 10% SDS PAGE followed by Coomassie staining. Pre-stained molecular weight markers were obtained from Thermo Scientific, USA. Protein concentrations were checked at every step with the help of Bradford reagent (Bio-Rad, USA) in a UV/visible spectrophotometer (The SmartSpec 3000, Bio-Rad, USA).

### Western blot

5−10 μg of proteins were separated on 10% SDS-PAGE and were consecutively electro blotted onto 0.45 µm nitrocellulose membrane (Millipore, USA). The membrane was blocked with 5% non-fat skim milk in PBS-0.1% Tween 20 for 2 hours at room temperature. After washing thrice with PBS-0.1% Tween 20 the membrane was probed with primary antibody which is a rabbit monoclonal anti human lamin A+C antibody (Thermo Scientific, USA) and Stabilized Peroxidase Conjugated secondary Goat Anti-Rabbit (H+L) antibody (Thermo Scientific, USA) at dilutions of (1:200) and (1:3000) respectively. The blots were developed by chemiluminescence using Supersignal Westpico Chemiluminescent substrate (Thermo Scientific, USA) and developed on Kodak Medical X-ray Films as described earlier [27]. All the blots were repeated at least thrice for consistent results..

### Circular Dichroism spectroscopy

Far-UV CD spectra of 2 μM human lamin A purified and degassed proteins (wt LA and A350P) were analysed using a Jasco J-815 spectropolarimeter (JASCO International,USA) to check the secondary structure of the proteins after dialysis from the denatured state in 6 M urea. A cuvette having 0.1 mm optical path length and a bandwidth of 1 nm was used. Solvent spectra recorded from working buffer were subtracted from the measured spectra in each experimental set. All spectra were recorded in the far UV range of 200–250 nm at 25°C, at a scan rate of 10 nm/min in a continuous mode considering an average of ten scans. Spectral analysis and smoothening was performed by Savitzky-Golay algorithm using Origin 8 Pro software. Data in triplicate was obtained for each protein.

### Steady state fluorescence spectroscopy

Steady state fluorescence spectra of renatured full length LA proteins (wt LA and A350P) in working buffer were recorded in a Hitachi F7000 spectrofluorimeter (Hitach, Japan). A quartz cuvette of 1 cm path-length was used for the spectra measurement of 4μM proteins at 25°C keeping excitation and emission slits at a band pass of 2.5 nm to obtain a good signal-to-noise ratio. Background intensities were omitted from each sample to cancel out any contribution due to the solvent Raman peak and other scattering artifacts. Since N-acetyl-L-tryptophanamide [NATA (Sigma Aldrich, USA)] dissolved in the working buffer works as a model for completely exposed tryptophan residues, all spectra were normalized to NATA. Correction for inner filter effect wasn’t necessary as the absorbance values of proteins at 340 nm were in the range of 0.03 and 0.01. Replicates of 3 in each set were achieved for all fluorescence spectra and individual spectral analysis was accomplished with the Origin 8 Pro software.

### Optical tweezer based microrheology

The schematic of experimental setup is given in Supplementary Figure 1. The optical tweezer was built using an inverted microscope (Olympus IX71, Japan) with 100x 1.3 numerical aperture (NA) objective for tightly focusing Gaussian laser beam of wavelength 1064 nm. A semiconductor laser (Laserver, maximum power 500 □ mW) was used to create a potential minimum to trap the particle inside the protein sample. The sample was prepared out of a small volume fraction (*ϕ* ≈ 0.01) of polystyrene particles of diameter 3 μm in a fixed concentration of protein, around 20 μl of which was drop-cast on a standard glass coverslip of thickness 160 μm and placed on top of the objective lens. A closed chamber was prepared around the sample so that no external perturbations (mechanical, electrical, or thermal) could affect the experiment. Indeed, the entire setup was mounted on a vibration isolation optical table so as to minimize mechanical contact with the external world. The modulation to the trapped polystyrene particle was provided using an acousto-optic-modulator (AOM of Brimrose, USA) which produced two beams at its output after the trapping laser was coupled into it – a spatially and frequency-wise unshifted zeroth order beam and a spatially and frequency shifted first order diffracted beam. This first order diffracted beam was modulated by a square wave pulse of frequency 1 Hz and amplitude 120 nm, and coupled into the sample space. A second 780 nm laser was also focused on the probe to detect its displacement using a balanced detection system [34]. The particle response was measured using two photodiodes interfaced with a data acquisition system (DAQ) at 5 KHz, with an acquisition time of 200 s. The Brownian motion of the probe without modulation, which was required for trap calibration, was captured at 10 KHz for an acquisition time of 20 s. It was ensured that the particle was trapped away (7 times of the particle diameter) from the coverslip to avoid the surface effect [35]. The response of the trapped Brownian probe is governed by a general Langevin equation which assumes a stationary and isotropic fluid around the particle, so that the equation can be written as,

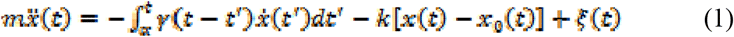

where, the mass of the particle is *m, k* is the stiffness of the harmonic potential, and *x*_0_(*t*) is the instantaneous displacement of the particle near the minimum of the potential. The integral term on the RHS is generalized time-dependent memory kernel which is associated to the damping of the fluid. The last term of Eq.1 represents the Gaussian distributed correlated random force applied by the surrounding fluid molecules which acts on the particle at every possible frequency. This thermal noise term can be written as, < *ξ*(*t*) *ξ* (*t* ′) > = 2*k*_*B*_ *T γ* (*t* − *t* ′) where *k*_*B*_ is the Boltzmann constant and T is the temperature of the system. Since the system is over-damped, the inertial term here can be ignored and the Eq.1 can be written in Fourier domain as,

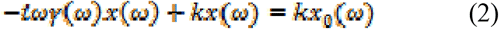

The frequency dependent friction coefficient is complex in nature for the viscoelastic fluid and it can be decomposed as, *γ* (*ω*) = *γ* ′ (*ω*) +*t γ* ″ (*ω*), and the phase response of the particle is,

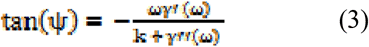

Here, the negative sign implies that the phase response lags the external perturbation (*kx*_0_) to the system. To extract the desired parameter values of viscoelastic property two different values of *k* i.e. *k*_1_ and *k*_2_ were required from which two different values of, *ψ* say *ψ*_*1*_ and *ψ*_2_ could be calculated. Therefore, Eq.3 can be rewritten using Eqs. (2) and (3) provided in the previous section, so that we have,

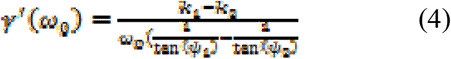

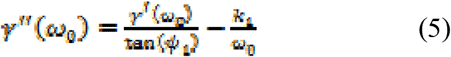

For a linear viscoelastic fluid, the relation between complex shear modulus and friction coefficient is *G* ″(*ω*) = − *t ω γ*/6 *π η*_0_, where *η*_0_ is the bulk viscosity of the fluid. Once again, complex modulus could be simplified into real and imaginary parts, which relate the loss moduli (*G* ″) and storage moduli (*G* ′) respectively, with the other parameters as,

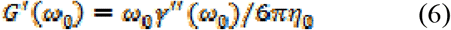

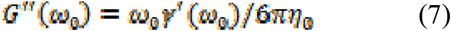

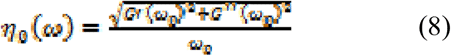

### Cell culture and transfection

Human Skeletal Myoblast cells (Gibco,Thermo Scientific, USA) were grown in DMEM (Gibco,Thermo Scientific, USA) media containing glucose to a final concentration of 10mM, supplemented with penicillin-streptomycin to a final concentration of 1% and Fetal Bovine Serum (Gibco,Thermo Scientific, USA) to a final concentration of 10%. The cells were maintained in a humidified incubator at 37°C and 5% CO_2_. When the cells grown on cover slips reached 50 to 60% confluency in serum free media, they were transfected with EGFP tagged wt LA/A350P DNA along with Lipofectamine 2000 (Invitrogen, USA) in a ratio 1:1.5. Serum complemented media was added after 6-8 hours and after this the cells were maintained at 37°C for 24 hours. The cells were subsequently fixed in 4% paraformaldehyde for 15 minutes and the coverslips were mounted with Vectashield + DAPI (Vector Laboratories, USA) on a glass slide. Myoblast cells which were positively transfected with EGFP tagged wild type LA/ mutant LA were used for all the *ex-vivo* experiments.

### Image acquisition, analysis and statistics

A NIKON Inverted Research ECLIPSE TiE Laser Scanning Confocal/NSIM Microscope (Nikon, Japan) was used to capture images of the transfected human myoblast cells. For the confocal microscopy, Plan-Apochromat VC 100 x oil DIC N2/1.40 N.A/1.515 RI objective was used and images were captured with a NikonA1R photomultiplier detector (Galvano mode). For the green channel, the lasers used were a solid state diode laser (λex = 488) used at 5% of the original wattage (50 mW) and for the blue channel a solid state diode laser (λex =405 nm) was used at 6% of the original wattage (100 mW). An additional 4x digital zoom was used to visualize the nuclei. A super resolution Plan Apochromat TIRF 100x/1.5 NA/WD 0.13 mm (Nikon, Japan) objective was used in the confocal mode to capture the pattern of lamin A meshwork formed in the cell nuclei by wild type and mutated proteins. NSIM-multi line laser (Coherent Sapphire 488 nm) was kept at 15% of the original wattage (200 mW) for the experiment. The 3D deconvolution of the final image from 15 images was carried out with the help of 3D blind deconvolution module of Autoquant ×2 (Media Cybernetics, USA). The inbuilt NIS-Elements acquisition software Ver.4.13.00 which operated under a windows environment was used to acquire and save the images of the nucleus as well as to control the elements both in the confocal and in the super resolution mode. The images were further analysed with the Image J software (1.46r) and mesh area was calculated from the Measure tool under the Analyze tab by assuming the mesh outline was roughly an ellipse. The data was exported to excel (Microsoft Excel 2010) and analysed for statistics.

## Results

### Expression of full length heterologous protein

The position of the missense mutation in lamin A protein backbone is shown in Figure 1A and B. The proteins were overexpressed in a heterologous system by inducing 2xYT medium with 1 mM IPTG. Both wt LA and A350P proteins were purified by using an anion exchange column and a size exclusion column sequentially to near homogeneity (>95% purity). Thus the residual impurities persisting from anion exchange column were completely removed by Size Exclusion Chromatography (SEC). In terms of recovery of proteins after denaturation by 6 M urea, wt LA showed fewer propensities for aggregation while dialyzing out urea than A350P. 10 µg of proteins were loaded for SDS PAGE analysis (Figure 1C) while 5 µg of proteins were loaded for authentication by western blot (Figure 1D).

### Analysis of secondary structure of the proteins by CD spectroscopy

Figure 2A shows the far-UV CD spectra of the wt LA and A350P at 2 µM concentration thereby validating the fact that the proteins were correctly refolded and predominantly α-helical as reported earlier and is in good agreement with the crystal structure [14, 28, 36]. Apart from the confirmation of a positive renaturation of the two proteins, the comparative analysis of the spectra also attests to modified secondary structural properties as reflected by the mutant with a probable change in the backbone structure. At a similar concentration, the two proteins exhibited non-overlapping spectra indicative of alteration in the secondary structure in A350P. The ratio of the ellipticity values at 222 nm and 208 nm (Figure 2A [inset]) was clearly higher for wt LA. Even though definitive structural predictions cannot be made from the given data, the decreased ratio in A350P could be attributed to lower helicity as [□]_208_ determines the percentage of α-helix [36]. Chromophores other than amides have minimal effect in the 208-230 nm regions so any dissimilarity observed could originate from the nature of helix of the respective proteins.

**Figure 2.**
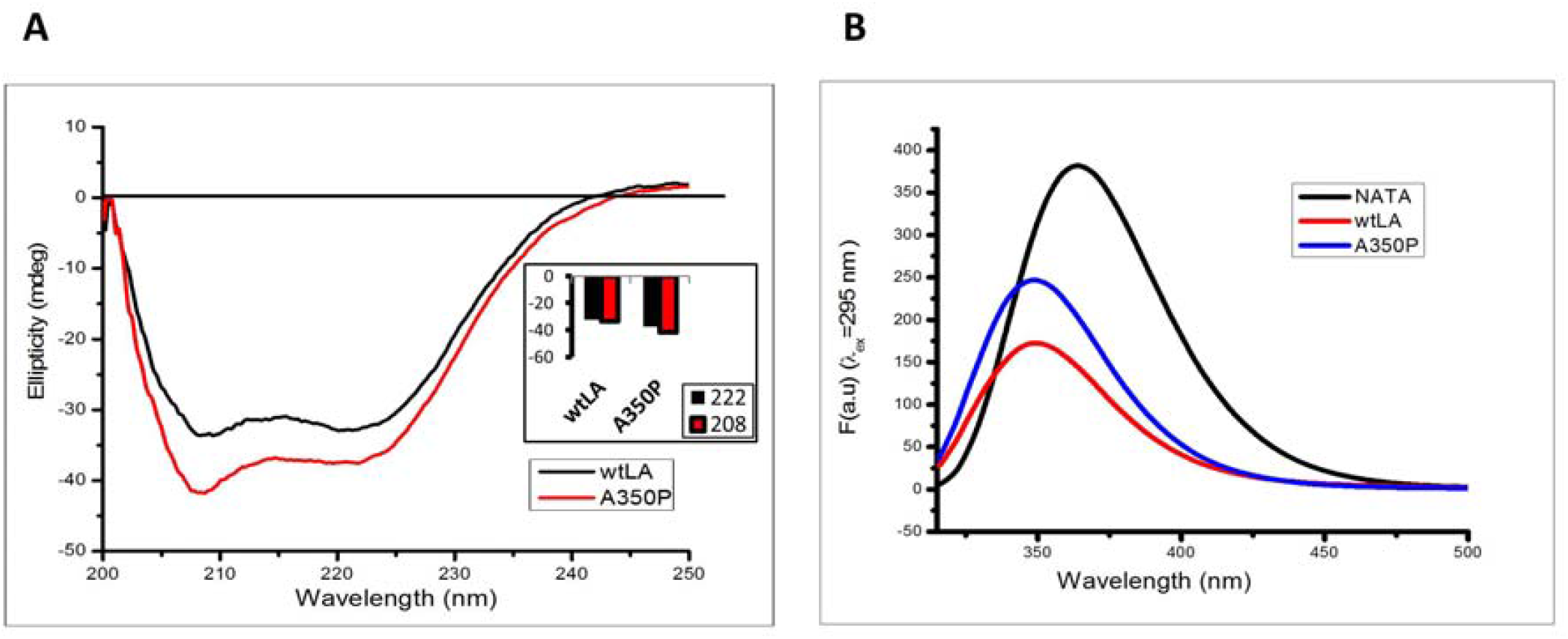
Secondary and tertiary structure of lamin A proteins: (A) Secondary structure of lamin A is altered by point mutation. CD spectra of 2 μM wt LA and A350P and comparison of observed values at 222 nm and 208 nm for the wt LA and A350P proteins (inset). (B) Mutation perturbs the tertiary structure of LA. Fluorescence spectra (λex= 295 nm) of 4 μM LA proteins for tryptophan residues and 16 μM NATA.

### Fluorescence spectroscopic studies for tertiary structure determination

As a sequel to the CD spectroscopic studies, we recorded steady state fluorescence measurements for both wt LA and A350P at 4 µM concentration to investigate any plausible changes in the tertiary structure. The spectra of the full length proteins were normalized using N-acetyl-L-tryptophanamide (NATA) where the tryptophan residue is fully solvent exposed. A distinct blue shift was noted in the spectra for both the proteins relative to NATA (Figure 2B) demonstrating the hydrophobic environment of the Trp residues referred to above. Though emission maxima did not vary much between the proteins but A350P showed higher quantum yield than wt LA at the same concentration. This might point to a slightly more compact hydrophobic core in the mutant compared to the wild type.

### Derivation of elastic moduli from optical tweezer based active microrheology

In order to measure viscoelastic parameters of the lamins, we needed to first calibrate our optical tweezers. This was performed by trapping a Brownian particle (radius 1.5 μm) and measuring its Brownian motion to determine the trap stiffness of our tweezers using the Equipartition theorem. The trap stiffness is defined as, 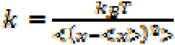, where < (*x* − < *x* >)^2^> is the variance of Brownian noise of the trapped particle. We modulated the trapped particle using a square wave, and checked the phase response, which should follow, tan^-1^(*f* / *f*_*c*_) where *f*_*c*_ denotes the corner frequency which is analogous to the trap stiffness (*k* = 12 *π*^2^ *η α f*_*c*_). For two sets of laser powers, we obtained corner frequencies of 116.6 ± 7.4 Hz and 231.1 ± 20.0 Hz, which led to trap stiffness of 17.61 ± 1.12 *μ* N/m and 34.87 ± 3.02 *μ* N/m, respectively. We also verified that the trap stiffness varied linearly with laser power. Subsequently, we used a modulation frequency 1 Hz and amplitude of 110 nm for the square wave in order to measure rheological properties of lamin A proteins by active microrheology. The advantage of applying a square pulse was to incorporate all the odd harmonics of the fundamental frequency in this signal by construction – being directly demonstrable by a Fourier domain decomposition. Mathematically, this may be represented as,

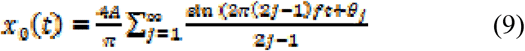

where, the displacement is *x*_*0*_*(t)*, ***A*** is the given amplitude, *f* is the fundamental frequency of the modulation, and *θ*_*j*_ is the modulation phase of the *j*^th^ harmonic. We used lock-in detection (*20*) to extract the phase response of the particle at each frequency for two trap stiffness values. From theory (provided in the Materials and Methods section), we have

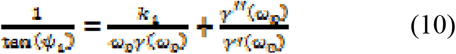

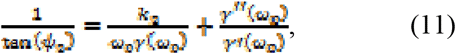

where *k*_1_ and *k*_2_ are the trap stiffness at two powers, giving rise to phase responses of *ψ*_1_ and *ψ*_2_, respectively, and *ω*_0_ is the modulation frequency, and *γ*(*ω*) = *γ*′(*ω*) + *tγ*″(*ω*), where γ(ω), is the complex friction coefficient, *γ*′(*ω*) with being the real component, and *γ*″(*ω*) the imaginary component of the friction coefficient. Solving the Eq. (2) and (3), we obtained *γ*’ (*ω*_0_) and *γ* ″ (*ω*_0_), which further led us to calculate the loss moduli (*G* ″) and storage moduli (*G* ′) following the eq. (6) and (7).

The results of our experiments are depicted in Figure 3A and 3B. We determined the storage moduli, (*G* ′) and loss moduli (*G*″) over an angular frequency range of 6-1100 rad/s – which is typically much higher than that accessible in commercial rheometers that are typically used for passive microrheology. We used three concentrations of both wt LA and A350P: 0.5 mg/ml, 1 mg/ml and 1.5 mg/ml. At higher concentrations, the protein didn’t form stable dispersions with water and we observed coagulation in different regions of the sample. Both (*G* ′) and (*G* ″) increased with increasing frequency which is consistent with the theoretical understanding of complex viscoelasticity as a function of frequency. Storage moduli (*G* ′) for the mutant A350P lamin was consistently higher than that for wt LA by a factor between 2.5-3 as is evident from Figure 3A. To clearly visualize the significant increase in the storage modulus, we explicitly plotted the ratio of the mutant type to the wild type at the same frequency for all the concentrations in Figure 3C. Thus, the increase in the storage modulus is more for higher concentrations and low frequencies where we see even an order of magnitude change for the mutant compared to wt. It is clear that the complex viscosity for A350P is higher than that of the wt LA over the entire frequency range that we measured. But at higher frequencies, the difference between the mutant and the wild type was reduced for the highest concentrations of the proteins due to the fact that (*G* ″)became higher for the wild type at those frequencies.

**Figure 3.**
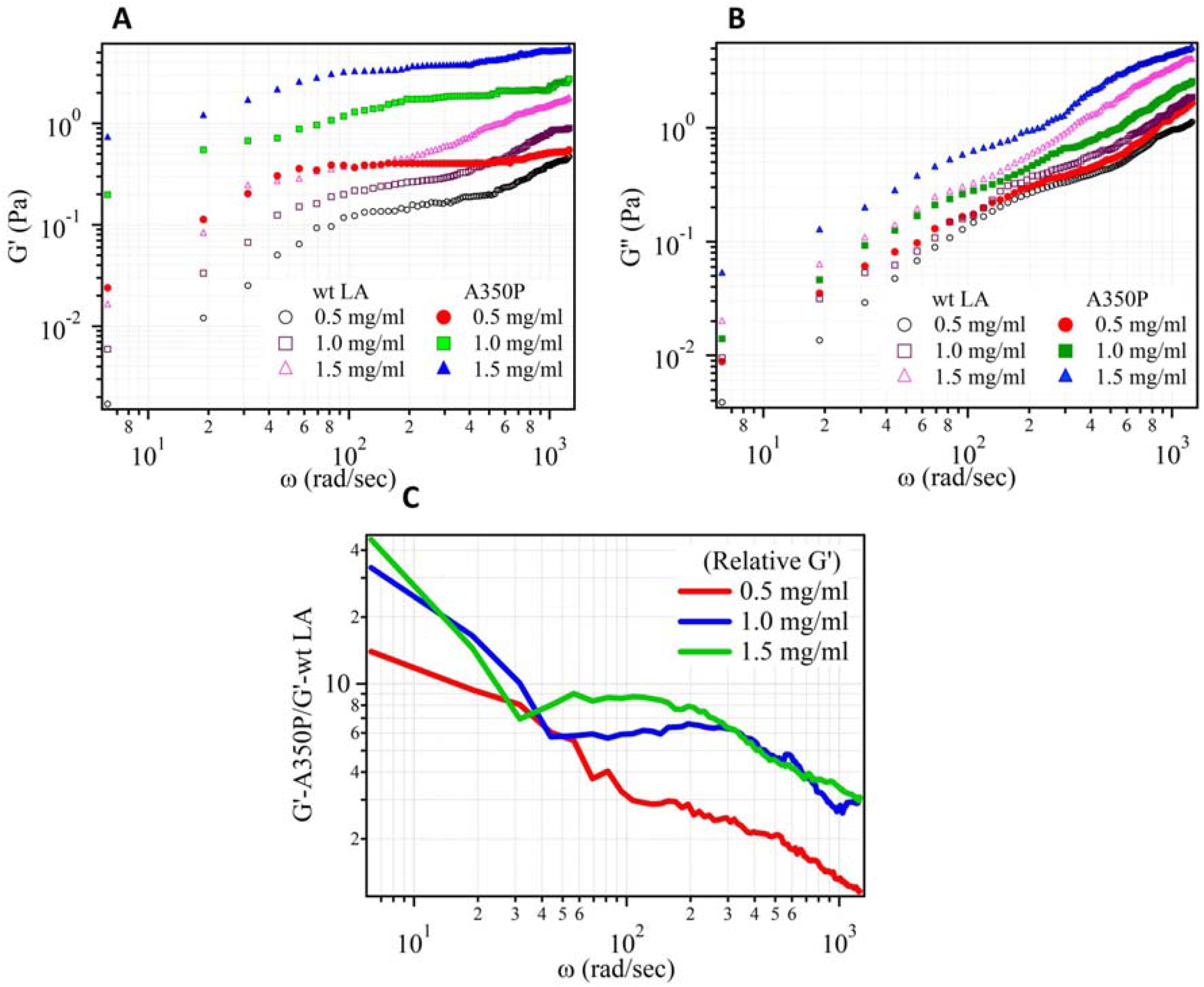
Comparison of (A) storage moduli (G’) and (B) loss moduli (G’’) between wt LA and A350P with frequency and varying concentrations of the proteins.(C) Relative storage moduli of A350P with respect to wt LA

### Analysis of mesh architecture by super resolution microscopy

While the secondary and tertiary structural alteration in the lamin A protein as induced by the single amino acid mutation hints on great physiological relevance – it cannot effectively explain the increased value of G’ for A350P. Hence, our subsequent experiment was designed to provide a rational explanation for the altered viscoelastic parameters in the mutant protein. When human myoblast cells were transfected with EGFP tagged wt LA /A350P, the wild type LA showed an intact rim of lamin protein along the nuclear lamina and a sparsely distributed continuous mesh of the protein prevailing throughout the nucleoplasm which was similar to previous observations [16, 21, 27]. Interestingly in case of the mutant, the bright rim was visibly absent. Instead, proteins formed in most cases isolated big denser network islands in the nucleoplasm in a highly random fashion (Figure 4A). This is a completely new observation of the phenotype from the networks of a lamin A mutant inside the nucleus. Super resolution microscopy revealed the fine difference in the nature of the networks thus formed by the proteins (Figure 4B) as analysed by mesh area calculation. Average mesh area for wt LA was observed as 0.0043 μm^2^ while for A350P it was 0.003 μm^2^. For A350P, the average mesh area was almost 43% lower than the wt LA (Figure 4C) which was rigorously calculated for 100 mesh structures, each from 10 individual nuclei. This could mean that even though the nuclear lamina remains vulnerable to mechanical stress because of the absent rim of protein, the A350P mutant could give rise to a much stiffer network inside the nucleus, which is essentially nonfunctional in nature.

**Figure 4.**
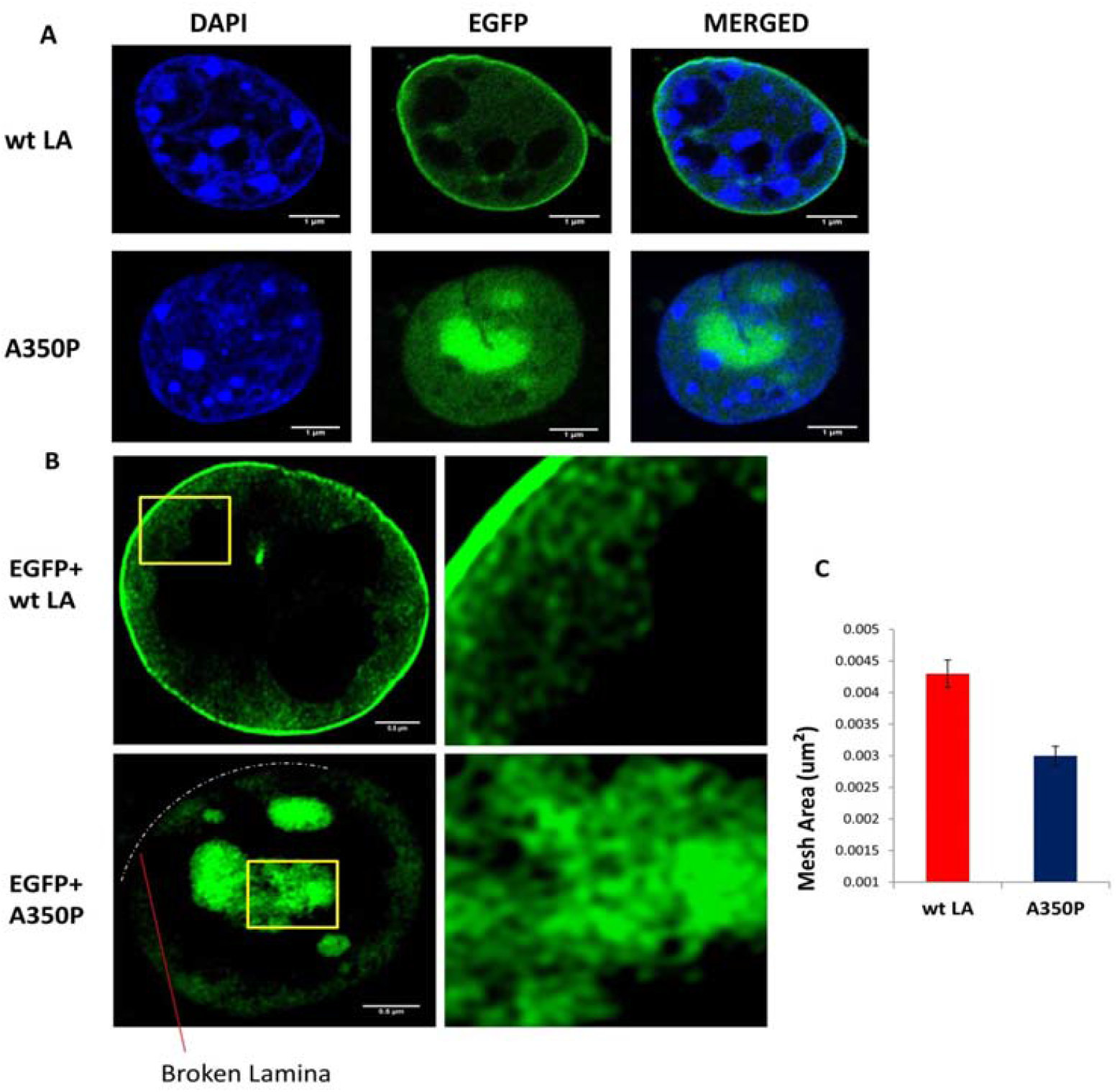
The network patterns formed by lamin A and its mutant in a human myoblast cell. (A) The different types of meshwork formed by wt/mutant LA in human muscle cell lines. The panel named DAPI shows the nuclear content of the cell at 405 nm. The second panel shows confocal images of Human Myoblast cells expressing EGFP tagged wt LA and A350P mutant at 488 nm. The merged panel shows the overlay of EGFP and DAPI channels. Scale bar is 1 μm. (B) The actual networks of wt/mutant lamin inside the human myoblast cell as observed under super resolution microscopy. The broken lamina in case of A350P mutant is indicated by dotted lines. (C) A comparative analysis of the mesh area formed by EGFP tagged wt LA ad A350P in the nucleus showing the mutant forming much denser networks. Error bar indicates percentage error. Scale bar is 0.5 μm

## Discussion

As we have mentioned earlier – in recent years, optical tweezers mediated force spectroscopy and microrheology has become an indispensable tool to study the biophysics of filament proteins, and has proved itself as an alternative to force spectroscopy using atomic force microscopes – which also have issues while dealing with live biological entities. Force spectroscopy using optical tweezers have been employed to determine the elastic properties of the cellular cytoskeleton [37, 38] and the extracellular matrix [39] from a force versus displacement curve of the trapped probe over multiple length scales [40]. The derivative of the curve obtained in the linear region provides quantitative information of the elasticity of the medium (essentially the spring constant) in which the probe is embedded (cytoskeleton on extracellular matrix), and also describes the force range where linear response is obtained. However – there also exists an entirely different methodology for obtaining the elastic properties of such environments – which is by measuring their complex shear moduli over a range of frequencies [41]. Thus, the real and imaginary components of the shear moduli, G’(ω) and G’’(ω) respectively, denote the elastic and viscous counterparts of the medium response, which elicit the same information of the medium as does force spectroscopy – with an additional component – which is its dissipative nature. This is performed by measuring the effects of the medium on the motion of a colloidal probe embedded in it – either by measuring its Brownian motion (passive microrheology [39], or its response to a drive (active microrheology [40]. However, to the best of our knowledge, active microrheology has not yet been applied to nucleoskeletal networks, but with some initial efforts to determine the amplitudes of the probe response in cytoskeletal protein keratin as described by Neckermuss *et. al*., 2015 and Paust *et. al*., 2016 [42, 43], without any attempts being made to determine G’(ω) and G’’(ω). Our results present the first effort in this direction, and suggest a major improvement over conventional techniques in active microrheology, which employ sinusoidal wave at a certain frequency, so that the response over a large bandwidth requires significantly long measurement times. The technique we employ here is based on our results described in [44] by deploying a square pulse for modulating the trap center. We obtained considerably larger signal to noise ratios compared to passive rheology, without compromising the bandwidth, and with much smaller measurement times compared to conventional active rheology using single sine waves. Additionally, we used the phase response of the probe compared to the conventional method of tracking the probe’s displacement amplitude, because the former is less prone to sensitivity issues and external noise, as clearly demonstrated by Paul *et al [44]*. Thus, we were able to correlate the viscoelastic properties to the probe’s trajectory, and thereby obtained an understanding about the mechanical structure of the nucleoskeleton in the wild type and mutant proteins, as we now proceed to describe.

We definitely observed an increase in the real component *G* ′ *(ω)* for A350P over the wild type lamin A in our measurements of the complex viscoelasticity employing oscillating optical tweezers. The increase in the storage moduli implied a higher crisscrossing of the protein polymers which eventually led to a denser protein network in the case of the mutant protein. This naturally entailed a smaller mesh size for the mutant protein, which was subsequently validated in the super resolution microscopy images. The higher storage modulus also implies a higher mechanical rigidity or lowered liquid-like behavior, which was manifested in the lower values of *G* ″ *(ω)* that we observed for the mutant protein compared to the wild type. The higher mesh density implied lower elastic response of the A350P to implied stress that might even lead to catastrophic failure on the application of localized stresses. The increased mechanical rigidity of the mutant lamin A may indeed be the reason behind the atrial fibrillation – a signature of DCM – which, as we mentioned earlier, is one of the dominant laminopathies leading to myocardial disorders in humans. Thus, we believe that the increased magnitude of *G* ′*(ω)* might be indicative of enhanced rigidity of the cardiac tissue, which makes our method of quantifying this parameter as a function of frequency a useful diagnostic tool in identifying early onset of atrial fibrillation or other cardiac laminopathies. It is important to note that the *in vitro* polymerized lamin A proteins closely resembles the *in vivo* network inside the cell and are easily interchangeable as systems to study this strain hardening behavior [28]. However, a careful characterization and quantification of both *G*′ *(ω)* and *G*″ *(ω)* for tissues in various degrees of fibrillation is required to set up a calibration procedure that is essential for employing our procedure as an efficacious diagnostic tool. Indeed, one may also extract proteins from tissues associated with distinct laminopathies and measure their viscoelastic properties to identify the existence of particular patterns which may then be employed for actual diagnosis of unknown patient samples. We have started work in these directions and expect to report very interesting results in the near future.

Apart from laminopathies, we hereby extend this method as a generalized tool to study the elastic properties of any protein and how that might change in a disease scenario. In the light of the SARS-Cov2 (COVID-19) pandemic, our method clearly promises significant implications. Post mortem autopsy from COVID-19 patients has been dominated by ground-glass opacities in the lungs, while histological hallmark included alveolar vasodilation which involved changes in primarily three categories of proteins namely surfactant proteins, extracellular matrix-cell junction proteins and core extracellular matrix proteins [45]. Interestingly all these proteins are mechanoelastic in nature and play significant roles in mechanical force transduction from the extracellular matrix to the interior of the cell. Therefore viscoelastic investigations using pulsed optical tweezers can be employed for microrheology-based investigations on these proteins to understand their role in the altered lung histopathology which in turn might enhance therapeutic intervention of COVID-19. The robustness and beauty of the system is highlighted by the fact that a minimal amount of protein – which being easily extractable from tissues – would elicit a response containing information about the elastic moduli (Figure 5).

**Figure 5.**
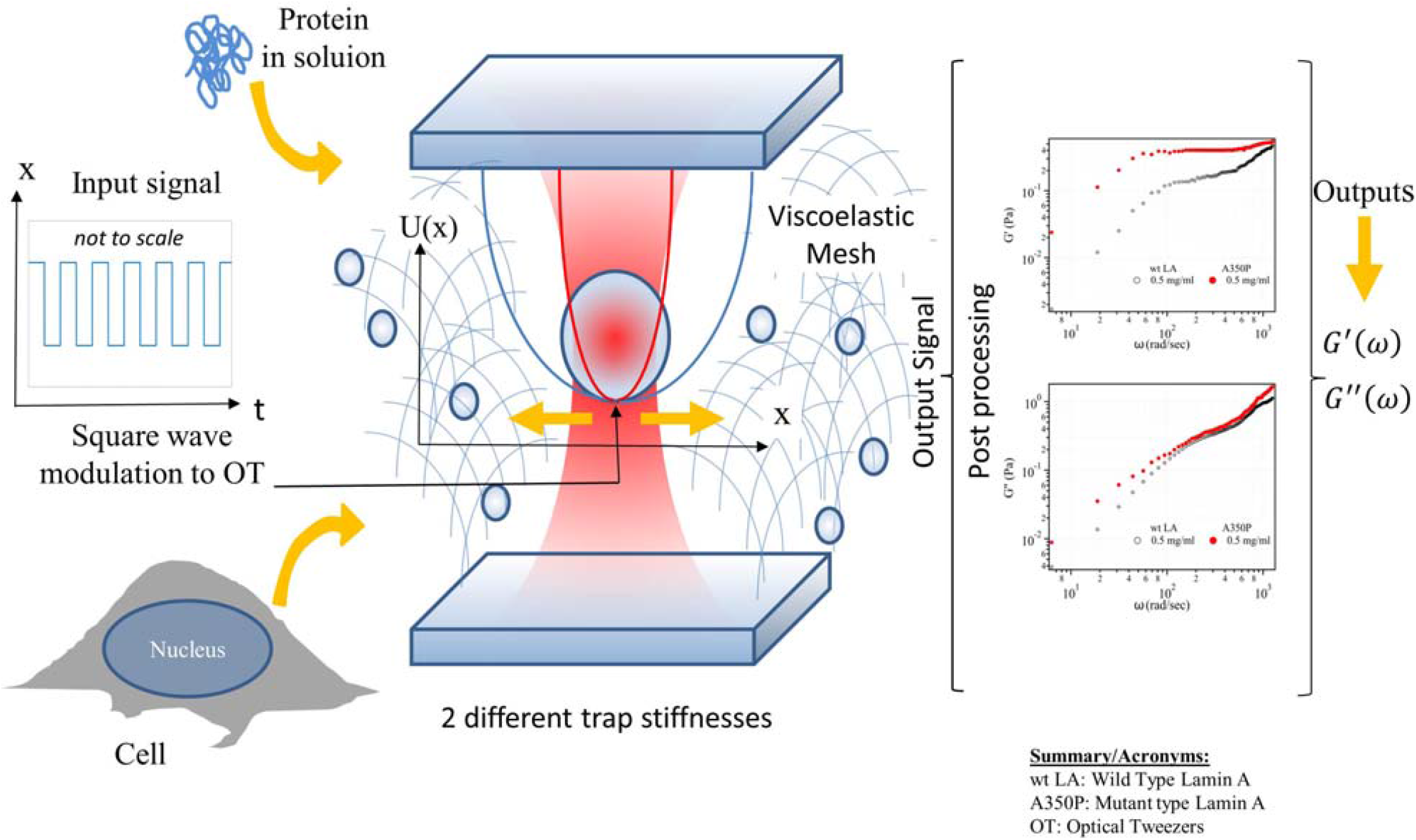
Derivation of G’ & G’’ from cellular or *in vitro* protein sample by pulsed optical tweezers.

**Figure 6.**
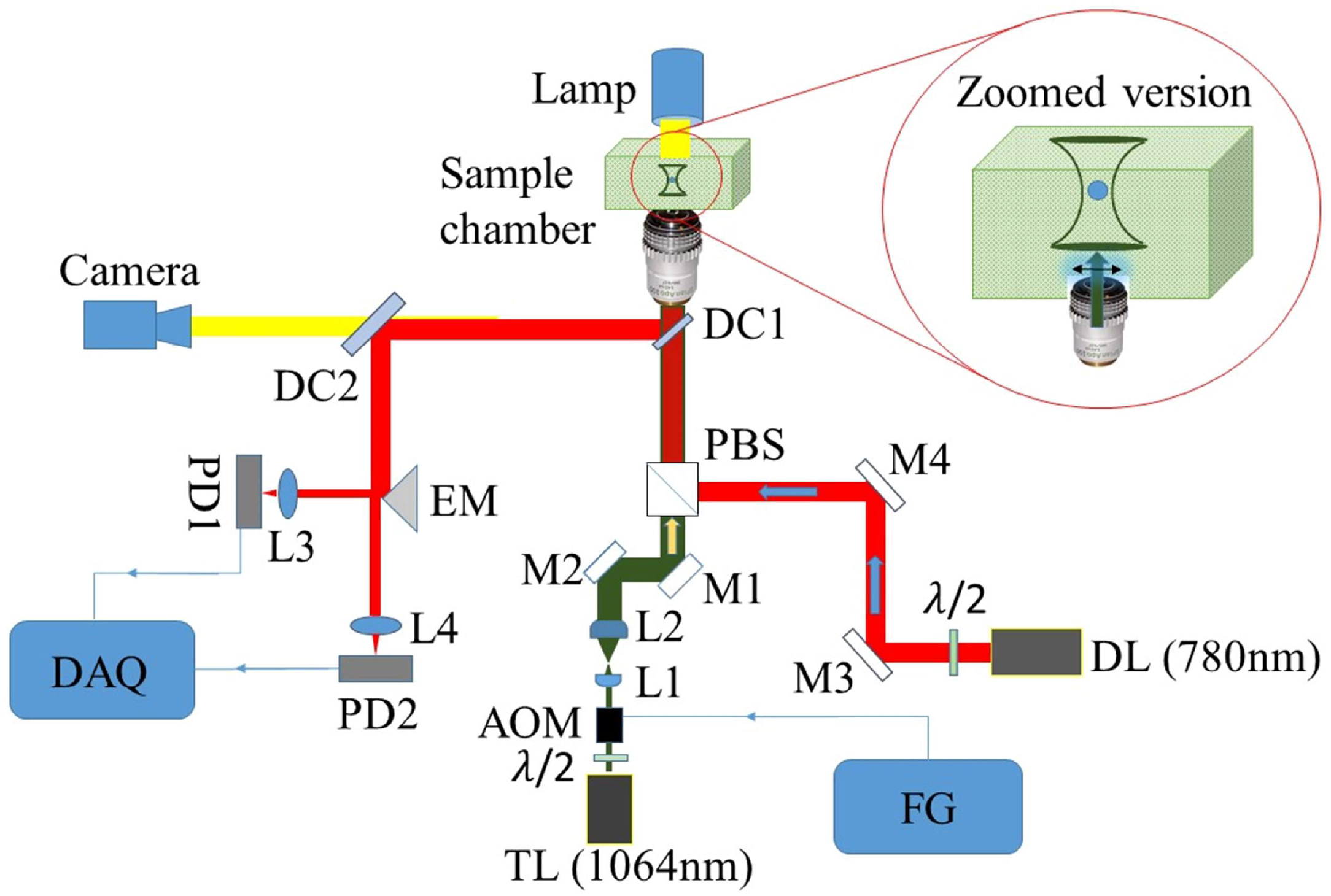

## Author Contribution

Chandrayee Mukherjee (CM), Avijit Kundu (AK), Raunak Dey (RD) performed the experiments, analysed the data and contributed in writing the paper. Ayan Banerjee (AB) and Kaushik Sengupta (KSG) conceived the project and wrote the manuscript.

## Acknowledgments

CM is funded by DAE, Govt. of India. AK and RD are funded by INSPIRE, DST, Govt. of India. AK, RD and AB also acknowledge IISER Kolkata. KSG is funded by BARD project, DAE, Govt. of India.

