## Supplemental Figure 1 for "Active microrheology using pulsed optical tweezers to probe viscoelasticity of Lamin A towards diagnosis of laminopathies"

**¶ Authors contributed equally**

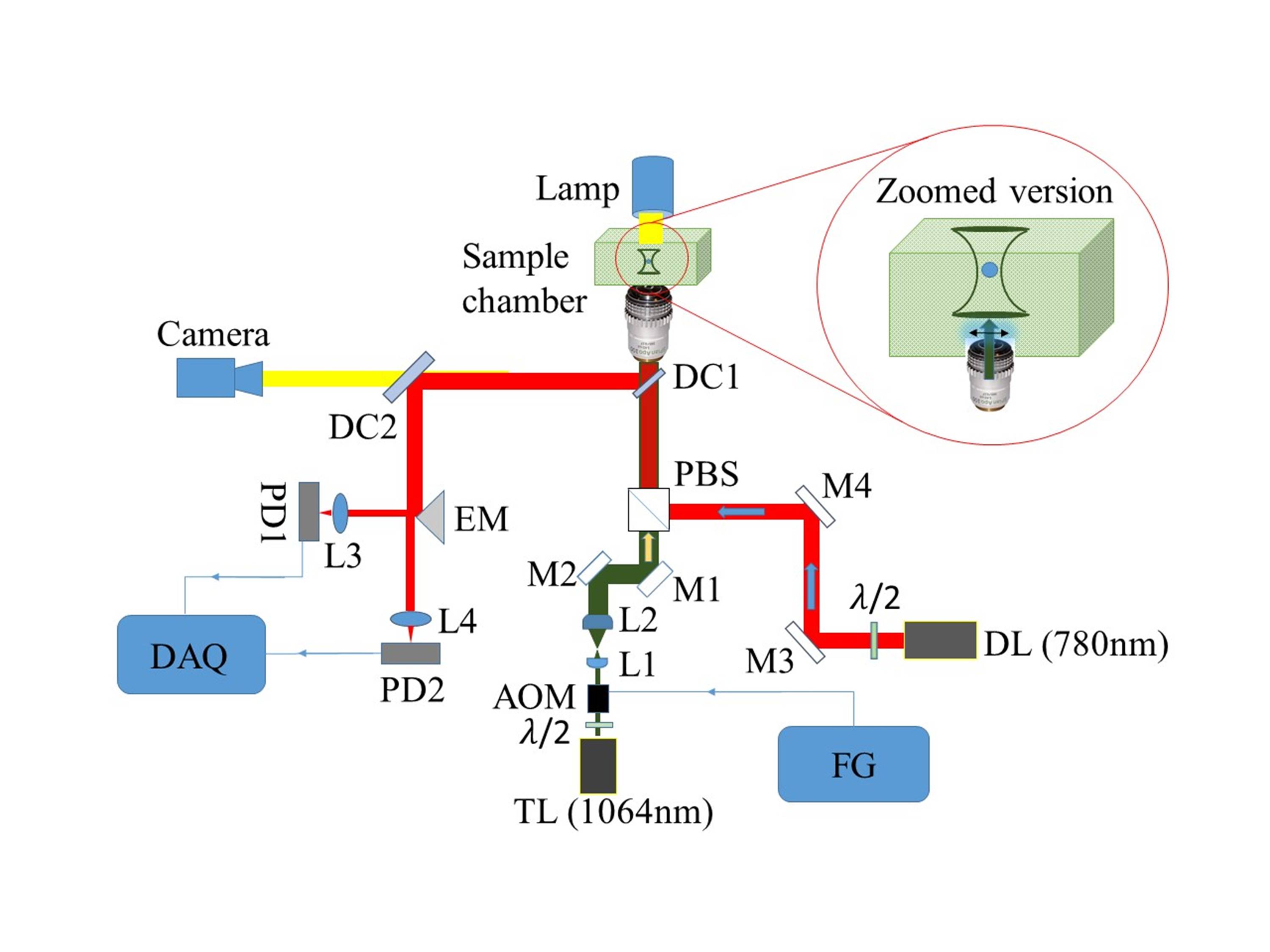

**Supplementary Figure 1. Experimental setup of pulsed optical tweezers**: Here TL: Traping laser ( λ = 1064 nm), DL: Detection laser ( λ = 780 nm), : Half waveplate, AOM: Acousto-optic-modulator, L: Lens, M: Mirror, EM: Edge mirror, DC: Dichroic, PBS: Polarizing beam splitter, PD: Photodiode, FG: Function generator, DAQ: Data acquisition card.
